# Causal dynamical modelling predicts novel regulatory genes of FOXP3 in human regulatory T cells

**DOI:** 10.1101/2020.02.13.943688

**Authors:** Rucha Sawlekar, Stefano Magni, Christophe Capelle, Alexandre Baron, Ni Zeng, Laurent Mombaerts, Zuogong Yue, Ye Yuan, Feng Q. He, Jorge Gonçalves

**Author notes:** Co-corresponding authors of this work.,. These authors contributed equally to this work (Rucha Sawlekar, Stefano Magni, Christophe Capelle).

## Abstract

Regulatory T cells (Tregs), characterized as a CD4+CD25+FOXP3+ subset of T cells, are vital to the induction of immune tolerance and the maintenance of immune homeostasis. While target genes of Treg master regulator FOXP3 have been identified, the upstream regulatory machinery of FOXP3 still remains largely unknown. Here we dynamically model *causal* relationships among genes from available time-series genome-scale datasets, to predict direct or indirect regulatory genes of FOXP3 in human primary Tregs. From the whole genome, we selected five top ranked candidates for further experimental validation. Following knockdown, three out of the five candidates indeed showed significant effects on the mRNA expression of FOXP3. Further experiments showed that one out of these three predicted candidates, namely nuclear receptor binding factor 2 (NRBF2), also affected FOXP3 protein expression. These results open new doors to identify potential new mechanisms of immune related diseases.

---

Regulatory T cells (Tregs) are key players of the immune system, which in turn plays a crucial role in a wide range of diseases. It is well established that the transcription factor FOXP3 is a master regulator of Tregs. However, genes regulating FOXP3 are still largely unknown and identifying them may be vital for developing new immunotherapeutics. This paper computationally screens the whole genome for potential regulators of FOXP3, and then experimentally validates them. Overall, the paper illustrates how the combination of genome-wide time-series data with dynamical modelling can identify a small set of relevant causal interactions.

Tregs perform immunosuppression of self-reactive lymphocytes to induce immunological self-tolerance and maintain homeostasis [Josefowicz et al., 2012; Kenneth et al., 2012; Li et al., 2015]. Tregs are involved in different types of diseases, such as autoimmune diseases [Dejaco et al., 2006; Fehérvari and Sakaguchi, 2004; Sakaguchi et al., 2006], cancer [Franchina et al., 2018; Shang et al., 2015; Tanaka and Sakaguchi, 2017], infectious diseases [Joosten and Ottenhoff, 2008; Stephen-Victor et al., 2017], neurodegenerative diseases [Baruch et al., 2015; He and Balling, 2013] and others [Cools et al., 2007].

The transcription factor FOXP3 has been shown to play a decisive role for the development and function of Tregs [Ziegler, 2006]. This gene is expressed specifically in CD4+CD25+ Tregs [Hori et al., 2003; Rudensky, 2011]. Altered expression of FOXP3 has been found in various types of autoimmune diseases [Liu et al., 2013] and tumors [Cunha et al., 2012; Martin et al., 2010; Szylberg et al., 2016]. Genetic mutations in FOXP3 result in autoimmunity and inflammatory syndromes, both in humans and in mice [Fontenot et al., 2003; Khattri et al., 2003; Mayer et al., 2014; Mercer and Unutmaz, 2009]. Most of the experimental evidence indicates that FOXP3 deficiency is responsible for IPEX (immunodysregulation polyendocrinopathy enteropathy X-linked) syndrome which is a rare disease caused by dysfunction of Tregs [Bennett et al., 2001].

Target genes of FOXP3 have been identified through intensive studies [Marson et al., 2007; Zheng et al., 2007; Zheng and Rudensky, 2007]. Meanwhile, progress in high-throughput technical developments, such as ChIP-seq and ChIP-Chip, have started to illuminate genetic and epigenetic mechanisms regulating expression or protein stability of FOXP3 [Chen et al., 2013; Floess et al., 2007; Fu et al., 2012; Gao et al., 2015; Miyara and Sakaguchi, 2007; Schmidt et al., 2012]. The majority of known upstream regulators of the expression of FOXP3 are general regulatory genes, e.g. those controlling interleukin signaling pathways (IL2, IL4, IL6 and so on) and cell surface receptors (TGFB) [Lal and Bromberg, 2009]. Those genes tend to regulate a large number of genes far beyond FOXP3, which might cause significant unwanted side effects when being targeted.

More specific upstream genes regulating FOXP3 are still largely unknown. Identifying these regulators of FOXP3 may be crucial for developing new immunotherapeutics against autoimmune and other related diseases. Unbiased experimental screening of the whole genome for such upstream regulators is almost impossible with current experimental approaches. However, such tasks can be efficiently performed computationally. This is the goal of this work.

The models trained in this work are based on microarray data of the whole-genome mRNA expression in isolated human Tregs [He et al., 2012]. Tregs from two different donors were stimulated at time zero with anti-CD3/-CD28/IL2, and measurements were taken at time zero followed by sampling every twenty minutes over a period of six hours (19 time points in total). After the pre-processing of the data, as outlined in column 1 of Fig. 1a and described in Supplementary Section 1.1, there were 13601 transcripts left, corresponding to 7826 genes. A gene can correspond to multiple transcripts, each measured by a separate probe set; from now on for simplicity we will only use the term transcripts and omit the term probe set. The role of the computational modelling was to reduce this number to 5 genes, reflecting our available experimental resources for validation.

**Figure 1:**
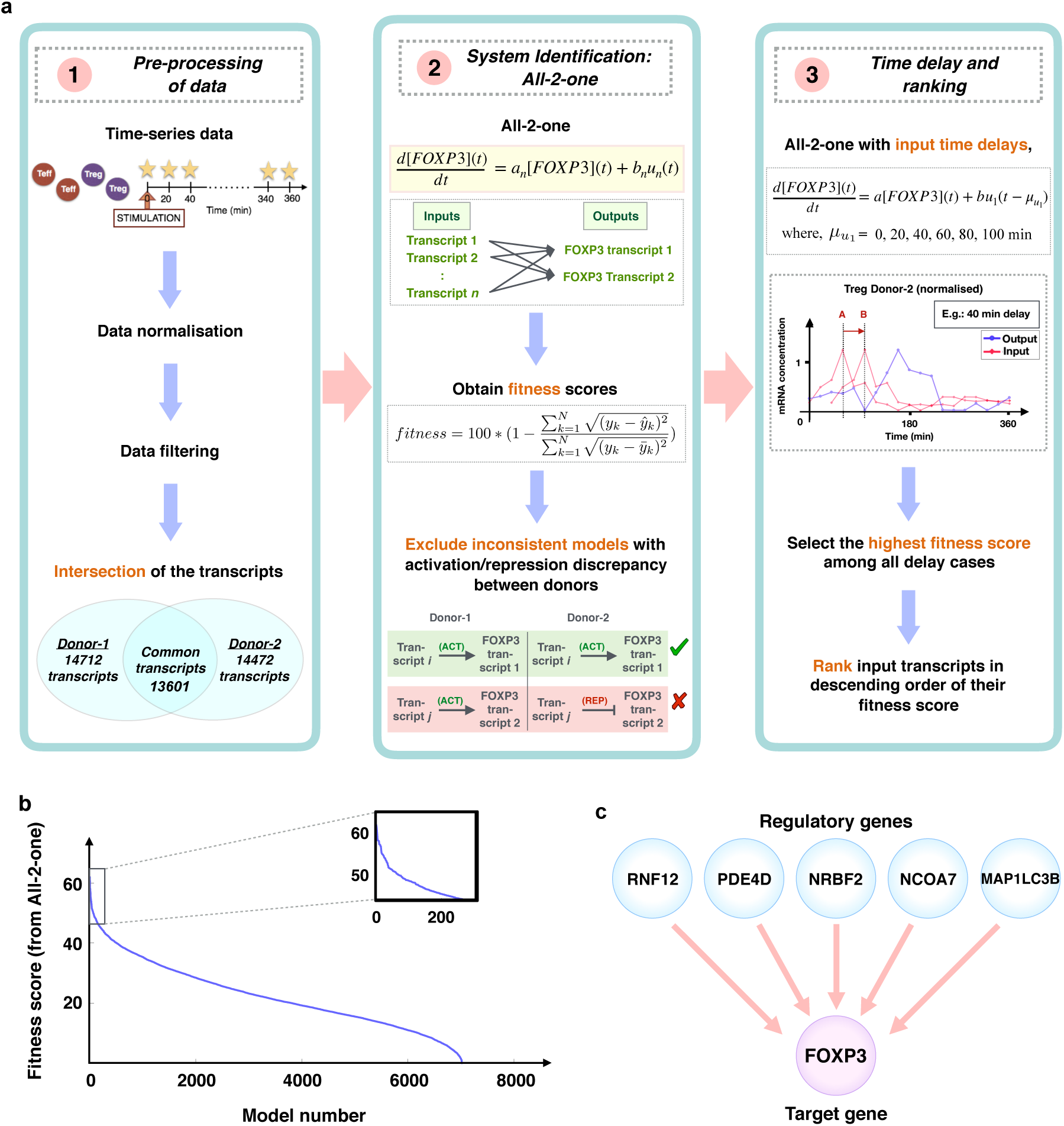
Causal dynamical modelling pipeline resulting in hypothesized regulators of FOXP3 for subsequent *in vitro* testing. **(a)** Overview of the different stages of the computational approach, from raw time-series transcriptomic data to the resulting ranking of genes likely to regulate FOXP3. The abbreviation ACT stands for activator, REP for repressor. **(b)** Fitness score versus model number for each transcript paired with each of the two available FOXP3 transcripts. Two regimes are visible: a group of models showing a steep decrease in fitness, and a much larger group showing an almost linear decrease in fitness. The box on the top right corner of this panel shows a magnification of the region with the top ranked models. The ranking of models directly translate in ranking the input transcripts. The rank of the first 176 transcripts is reported as Tab. S1. **(c)** Overview of predicted regulators of FOXP3 which were tested *in vitro*.

Existing models capture known regulations of genes by FOXP3 [Carbo et al., 2013, 2015; Hong et al., 2011; van den Ham and de Boer, 2008], so they were not useful in finding novel regulators of FOXP3. Hence, we need to fit new models from data that capture *causality*, i.e. that identify those genes that *cause* changes in FOXP3. Here, we take advantage of a relatively large number of available time-series Tregs samples to fit dynamical models. In particular, ordinary differential equations are mathematical abstractions that capture both dynamics and causality, and help build testable hypotheses in experiments. Our models take as inputs the time-series expression values of the whole genome, i.e. each of the 13601 transcripts left after pre-processing of the data, and FOXP3 as the single output (target).

There is always a trade-off on the choice of model complexity. With rich data, models can be complex, providing detailed information of mechanisms of action. However, with limited data, complex models can easily overfit and introduce bias, resulting in large numbers of false negatives. In our case, with only a single time-series experiment and limited resources for validation, we consider the simplest dynamical model: first-order linear time-invariant (LTI) systems, a well established modelling strategy [Dalchau, 2012; Herrero et al., 2012; Mombaerts et al., 2019b, 2016; Müller et al., 2019]. Moreover, to scale the method to the whole genome, we built a large number of simple pairwise models (one per transcript), testing whether each transcript on its own could regulate FOXP3. A model associated with a particular *transcript* is given by

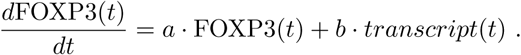

The left-hand side of the equation is the derivative of FOXP3 expression over time, i.e. the rate of change of the concentration of FOXP3. Finally, we searched for parameters *a, b* that best fitted the data. This procedure was repeated for all 13601 transcripts. Given the simplicity of the models, high fit means high confidence that such regulation of FOXP3 may indeed exist. This method was summarised in column 2 of Fig. 1a and further details are in Supplementary Sections 1.2 and 1.3.

Potential regulators of FOXP3 identified here may or may not be transcription factors. Indeed, our models can capture both direct or indirect regulations of FOXP3. Indirect regulations involve other molecules and typically require larger dimensional models to capture the dynamics of intermediate steps. To avoid increasing model complexity and still consider indirect regulations, we introduced a single time delay *τ* in the input signal

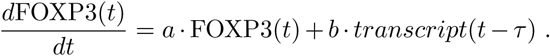

Hence, each model had a total of three parameters. Finally, transcripts were ranked according to how likely they are to regulate FOXP3. This was illustrated in column 3 of Fig. 1a and described in Supplementary Section 1.3.

We decided to focus on the top 176 transcripts (Tab. S1) out of 7030. Fig. 1b showed a significant change in the rate of change of the fitness values, i.e. a kink, after approximately the first 176 transcripts (additional details are provided in Supplementary Section 1.4). Of those, there were 161 distinct genes, since 15 of the 176 transcripts corresponded to the same genes. These genes covered fitness values ranging from 62 to 46 (the higher the fitness value, the better; 100 is perfect fit). Among the remaining 161 top ranked genes, FOXP3 is known to bind the promoters of 59 genes [Sadlon et al., 2010], and 38 genes are reported to be differentially expressed in human Tregs compared to CD4+CD25-effector T cells (Teffs) [Sadlon et al., 2010]. For 15 genes, both statements were true. As a first literature validation test, this shows that the predicted top ranked genes are indeed involved in the regulatory pathways related to FOXP3, already supporting the relevance of our prediction.

Next, we experimentally validated some of the 161 top ranked genes in primary human Tregs (Fig. 2a, details in Supplementary Section 2). Considering the available resources, we selected only 5 genes: *NCOA7, MAP1LC3B, NRBF2, PDE4D* and *RNF12* (Fig. 1c). The choice of these 5 genes was mostly based on a smooth dynamical profile with rise and fall dynamics ahead of FOXP3, and a diversity of activity in different cell parts. The details of this selection are provided in Supplementary Section 1.4.

**Figure 2:**
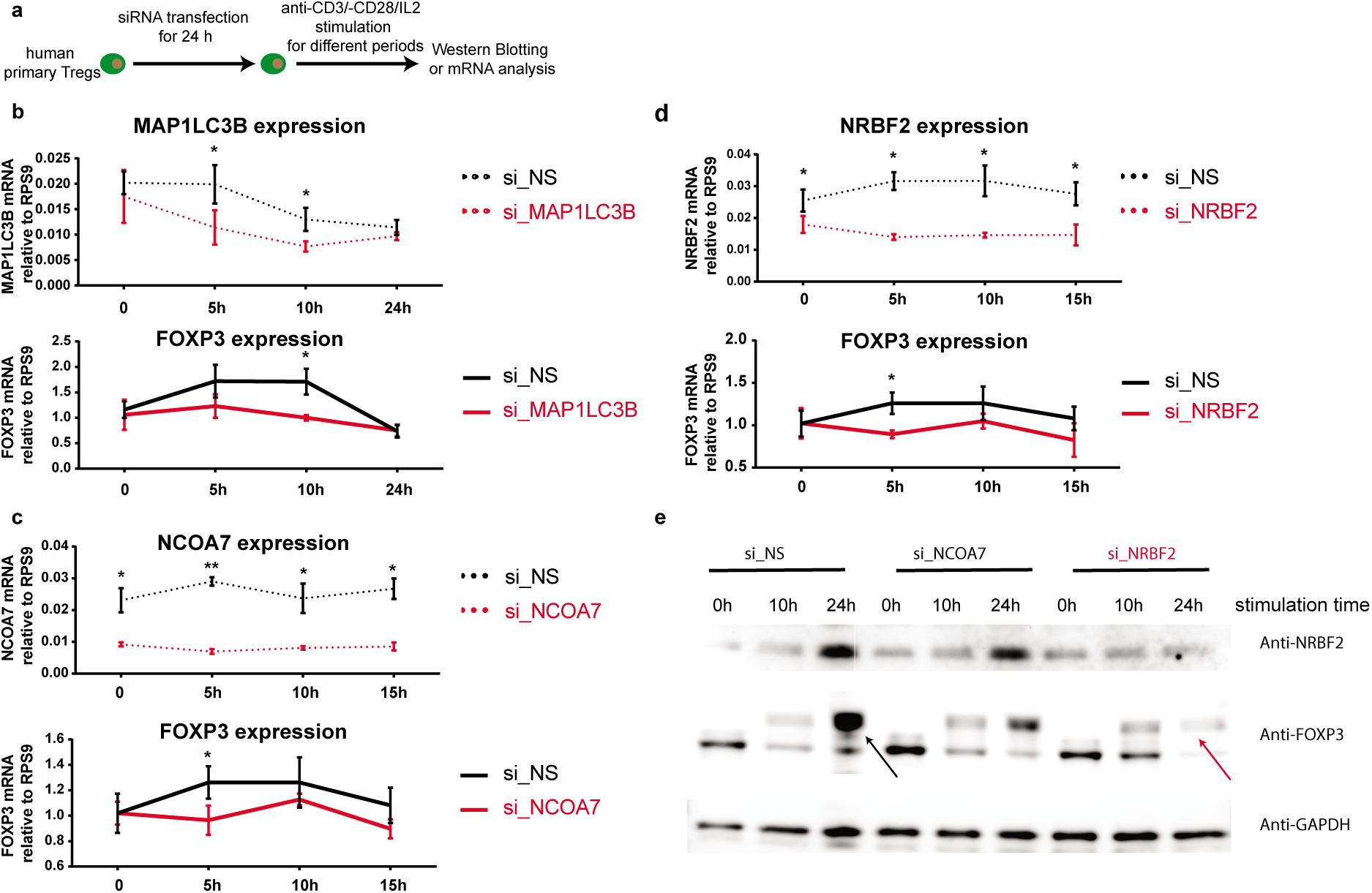
Experimental validation of predicted candidates. **(a)** Experimental scheme of the knockdown experiments. The predicted candidate genes were knocked-down by siRNA transfection in primary human Tregs for 24h, followed by CD3/CD28/human rIL-2 stimulation. **(b)** Quantitative real-time PCR results for the knockdown of MAP1LC3B in human primary Tregs and the corresponding FOXP3 expression. Control scrambled non-specific knockdown (si_NS) is shown in black and the specific knockdown (si_MAP1LC3B) in red. Statistical significance was determined using Student t-test, *p<0.05, **p<0.01 and ***p<0.001. **(c)** Quantitative real-time PCR results for the knockdown of NCOA7 in human primary Tregs and the corresponding FOXP3 expression. Control knockdown (si_NS) is shown in black and the specific knockdown (si_NCOA7) in red. **(d)** Quantitative real-time PCR results for the knockdown of NRBF2 in human primary Tregs and the corresponding FOXP3 expression. Control knockdown (si_NS) is shown in black and the specific knockdown (si_NRBF2) in red. **(e)** Western Blot showing the protein expression of NRBF2 and FOXP3 in human primary Tregs transfected with si_NCOA7, si_NRBF2 or control siRNA (si_NS). The bands of interest are highlighted by black (si_NS) or red arrows (si_NRBF2).

Our experimental results showed a successful knockdown of *MAP1LC3B, NCOA7* and *NRBF2* using siRNA specifically targeted against the corresponding gene relative to a control scrambled siRNA in primary human Tregs (Fig. 2b, c and d). Excitingly, knockdown of *MAP1LC3B, NCOA7* and *NRBF2* down-regulated the transcript expression of *FOXP3* in Tregs (Fig. 2b, c and d). The dynamics of *FOXP3* expression following siRNA treatment was slightly different for the three candidates. For the other two candidates, *PDE4D* and *RNF12*, although we successfully knocked down their mRNA expression, there was no clear effect on the mRNA expression of FOXP3 (data not shown here).

Next, we tested the potential effect on protein production. All our predictions were based on models built from mRNA data, not protein expression. Due to the limited availability of reliable antibodies on MAP1LC3B, we performed Western Blotting analysis to test the effects of knockdown of only NCOA7 and NRBF2 on the protein expression of FOXP3. So far, we could only successfully demonstrate the downregulation of NRBF2 protein expression. Notably, in line with the mRNA results, dowregulation of NRBF2 protein expression indeed reduced the expression of certain isoforms of the FOXP3 protein (Fig. 2e), possibly due to altered alternative splicing [Allan et al., 2005]. All these experiments have been successfully repeated in Tregs isolated from peripheral blood of 6 to 8 healthy donors, depending at which level (protein or mRNA or both) the validation was performed. The effect was not observed in 1 or 2 of the tested donors possibly due to the heterogeneous nature of human individual samples.

NRBF2 is known to regulate the activity of VPS34 [Lu et al., 2014]. Moreover, T-cell specific depletion of VPS34, significantly impaired the maintenance and the suppressor function of Tregs [Parekh et al., 2013]. Our results, together with these published works, already indicate that NRBF2 might be a promising upstream regulator that modulates the expression of FOXP3.

One limitation of our computational approach is that, being based on a linear dynamics, it might miss complex non-linear interactions. Furthermore, since our method checks one transcript at a time as a potential regulator of FOXP3, it might miss interactions requiring cooperativity among several regulators at a time. Due to limited amount of data available, however, higher model complexity is likely to lead to overfitting and false positives. Since the ground truth behind the biological system that generated this data is unknown, the safest route is to consider low complexity models, as we did here. Moreover, many transcription factors are regulated at the post-transcriptional level, which are not captured by our models and predictions (since the models are built on transcriptional data alone).

In summary, we applied dynamic modelling techniques to predict upstream regulatory genes of FOXP3, the master regulator of Tregs. Then, we experimentally validated the predicted effect of a handful of genes. Silencing of any of the three genes *MAP1LC3B, NCOA7* and *NRBF2* down-regulated the transcript expression of *FOXP3*. Moreover, dowregulation of NRBF2 protein expression reduced the expression of FOXP3 protein. Although NRBF2 knockout mice do not show spontaneous autoimmune phenotypes, NRBF2 positively regulates the autophagy process [Lu et al., 2014], which has been widely associated with autoimmune diseases [Gianchecchi et al., 2014]. Moreover, an integrative meta-analysis from around 72 million functional associations shows that NRBF2 is ranked as the sixth candidate gene related to juvenile rheumatoid arthritis [Rouillard et al., 2016], one of the classic autoimmune diseases. Overall, our results enhance our understanding of the upstream regulatory mechanisms of FOXP3, with the potential to develop new immunotherapeutics. Our results are solely derived from *human* primary T cells and therefore guarantee their application potential in medicine.

Potential future developments include, but are not limited to, the following: (1) to experimentally test *MAP1LC3B, NCOA7* and *NRBF2 in vivo* in animal models, for example using corresponding whole-body or T-cell-specific knockout mice under homeostatic and pathological conditions; (2) to experimentally investigate additional regulators of FOXP3 from the predicted list of top ranked genes; (3) to apply the same computational approach to identify regulators of CTLA4, another important gene of relevance for Tregs, and (4) to use more complex models capturing mechanistic details when, and if, new time-series data capturing different experimental conditions become available, including single-cell RNA-seq data [Chan et al., 2017] and [Ocone et al., 2015].

## Acknowledgements

The authors would like to thank Anthony Haynes, Frontinus, for his mentoring in writing and editing this manuscript. This work was supported by the Luxembourg National Research Fund (FNR) CORE grant, ref CORE/14/BM/8231540/GeDES. Stefano Magni and Christophe Capelle are supported in part by the FNR PRIDE DTU CriTiCS, ref 10907093. Jorge Gonçalves is partly supported by the 111 Project on Computational Intelligence and Intelligent Control, ref B18024. Feng He is partly supported by the Luxembourg Personalized Medicine Consortium (PMC), FNR individual AFR or AFR-RIKEN bilateral program ref 9989160 and TregBAR, PRIDE DTU NextImmune ref 11012546.

## SUPPLEMENTARY MATERIAL

### 1 COMPUTATIONAL METHODS DETAILS

#### 1.1 Time-Series Data Normalisation and Filtering

As shown in Fig. 1, first of all we obtained the raw data corresponding to Tregs cells from the online supplementary material of [He et al., 2012]. Therein, microarray measurements were performed for every 20 min over the period of 6 h, after stimulation by anti-CD3/CD28/human rIL2 at time 0 h on Treg cells from two donors (here referred to as donor-1, donor-2). This data contained 54676 transcripts/probe sets for each donor, mapping various transcript variants of almost each gene in the whole genome. For simplicity, we elsewhere only use the term transcript and skip the more technical term probe set, keeping in mind that they are in a one-to-one relation, thus exchangable.

Before applying any system identification technique, these time-series data need to be pre-processed. This involves normalising the data using the gcrma algorithm [Wu and Irizarry, 2010] which is implemented in MATLAB. For this and all the other computational aspects of this work, MATLAB versions R2016a, R2016b and R2017a were used. The normalised data were then subject to filtering. Firstly, we applied affymetrix flag filter, where any transcript was removed if marked as *absent* in every measurement taken at each instant of time. Conversely, we kept all the transcripts for which at least one measurement taken at any instant of time was marked with *marginally present* or *present*. The second filter applied removed the transcripts for which the average intensity (of the mRNA expression, which depends on the normalisation used above) is <50, or the largest intensity among measurements performed at any time is <100 (in the arbitrary units used by the gcrma algorithm). After this filtering, 14712 transcripts were left for donor-1 and 14472 transcripts for donor-2. The intersection of these two ensembles led to a common set of 13601 transcripts left.

#### 1.2 One-2-one Method

Here, we used the methodology presented in [Mombaerts et al., 2019b] to identify candidates regulators for FOXP3. This modeling strategy uses Linear Time-Invariant (LTI) models to capture the dynamics describing the rate of change of the selected transcript with respect to another input transcript. A linear modelling paradigm offers advantages when data are scarce. In particular, although linear models do not provide detailed functioning of the whole network, they are capable of identifying regulatory interactions with a reliable degree of precision (see below). A LTI model can be generally represented by the following set of equations:

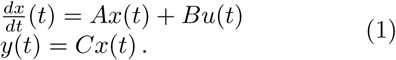

The model investigates whether the rate of change of the gene expression of a particular transcript *y*(*t*) depends on the gene expression of another transcript *u*(*t*). In particular, *u*(*t*) and *y*(*t*) represented the time series of the gene expression over time of a potential regulator of FOXP3 and FOXP3, respectively. The variable *x*(*t*) represented internal dynamics (translation, transcription, etc…) that interacted with the modelled output and were required for the behaviors observed, but were not explicitly included in the model. The dimension of the vector *x*(*t*) defines the model order: in general it can be a 1-dimensional vector (direct regulation or relatively slow dynamics compared to internal dynamics), or a multi-dimensional vector (the regulation happens through intermediate steps that introduce delays and cannot be ignored).

Estimating a model involves finding *A, B* and *C* which produce a vector *y*(*t*) as close as possible to the real expression data. On the one hand, complex nonlinear models have the potential to capture the dynamical relationships between genes with great precision. On the other hand, high complexity can lead to overfitting (fitting noise instead of dynamics) without sufficient data or detailed knowledge such as the network topology or types of nonlinear interactions. Here, we restricted the model order to one as it was optimally estimated in [Mombaerts et al., 2019b]. Hence, *A, B* and *C* were scalars. Furthermore, since *y* is the measurement of a single state, *C* then consists of a scalar of value 1. System identification was performed using the function ‘pem’ implemented in MATLAB to minimise the prediction error [Ljung, 2003].

Each model was characterized by a performance index that represents the capability of the model to describe the input-output relationship. To do so, we used the fitness:

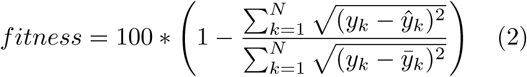

where *y*_*k*_ represented the data (output), 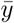 the average value of the data, and 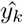 the estimated output. MATLAB function *compare* can be used to compute the fitness of the model. A fitness equal to 100% corresponds to a perfect identification. A high fitness suggests that most of the dynamics was captured.

Then, to investigate the potential regulators of FOXP3, a collection of 1st order LTI models was estimated from each of the transcripts to FOXP3. In each case, the parameters were estimated so that they together provided the best possible fit to FOXP3 time course data. This step took the following form:

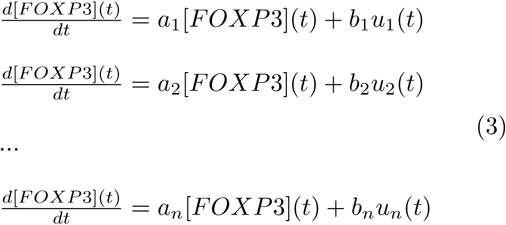

where *n* corresponds to the amount of candidates. Each model was characterized by a fitness metric that ranges from 0% to 100%, representing its capability of describing the original regulatory system between genes. A gene, therefore, would be further considered as a regulator for FOXP3 if the model obtained using one of its transcripts was capable of reproducing the expression profile of FOXP3 with a sufficient degree of precision, which should be characterized by a high goodness of fit, as defined above.

Additionally, we derived the mathematical expression of the dynamics of FOXP3 to explicitly include a delay in the modelling, such that it took the following form:

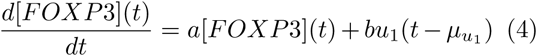

where 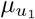 is a delay chosen between 0 min and 100 min, with steps of 20 minutes. The choice 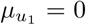 min reduces the models to the particular case employed above.

A systematic comparison of our methodology against state-of-the-art methods with data simulating similar experimental conditions can be found in [Mombaerts et al., 2019a].

#### 1.3 Applying the One-2-one to our data: All-2-one approach

For the system identification we only considered the transcripts that were left after filtering, as described above. Since, the number of transcripts differed between the two donors, we collected the common transcripts between the 14712 and 14472 transcripts (last step of column 1 of Fig. 1a). Our All-2-one algorithm used these remaining transcripts, one at a time, as an input (regulatory gene, represented by the variable *u*_*i*_ (*t*) in Eq. (3)), for the system identification technique described above as One-2-one (first step of column 2 of Fig. 1a). As output target, the two FOXP3 transcripts were used separately, one at a time. In fact, out of 3 measured transcripts of FOXP3, only two were considered here, because the third transcript associated to FOXP3 got discarded by the average intensity filter described earlier because of its very low expression. The All-2-one was repeated for each donor which gave us a total of 4 sets of All-2-one results (i.e. each input towards the two FOXP3 transcripts, for each of the two donors). The results contained fitness score of each input towards each output (second step of column 2 of Fig. 1a) and an indication whether the regulatory gene was tentatively an activator or a repressor of the target gene.

Out of the 4 sets of All-2-one results, we now combined the results of 2 FOXP3 transcripts within each donor (third step of column 2 of Fig. 1a). Then we discarded any input transcript which corresponded to two models (one in each donor) being inconsistent in showing activation or repression towards any of the FOXP3 transcripts (fourth step of column 2 of Fig. 1a). In fact, this would reflect an inconsistency between the inferred mathematical models. This yielded 3515 transcripts of inputs in each donor. Since there were two outputs, that meant 7030 models identified left (each with an associated fitness score) in each donor.

Typically, the time passing between the expression of a regulatory gene and that of its target gene is around 20 to 60 minutes. The One-2-one method tended to attribute higher fitness to models where the output signal follows the input signal pattern with one time-point distance, which in our case corresponded to 20 minutes. Thus, in order to identify regulations occurring over times longer than 20 minutes, we needed to modify the One-2-one method by introducing time-delays in the input signals only, Eq. (4). Thus, we ran a new round of All-2-one, with delays of 0, 20, 40, 60, 80 and 100 minutes, i.e. equivalent to moving each input signal to the right by 20-100 minutes (first step of column 3 of Fig. 1a). Here, for each input-output combination (i.e. model), only the highest fitness score among all 6 delay cases was retained (second step of column 3 of Fig. 1a). Even though for completeness this procedure was repeated exactly the same way for each donor, for the subsequent ranking we only utilised the results of donor-2, because in this donor FOXP3 mRNA expression showed a dynamics much more robust against noise. Eventually, these fitness scores were used to rank the 7030 models of donor-2 in a descending order (last step of column 3 of Fig. 1, see Tab. S1 in Supplementary Material for the high ranking part of the list of genes). This ranking was the main result of our computational method and higher the ranking of a gene, the more likely it is to be involved in FOXP3 regulation.

#### 1.4 Selection of Genes for Wet-Lab Experiments

The above mentioned results corresponded to hypothesizing several regulations, in particular the higher the ranking of a gene, the more likely it is to be involved in FOXP3 regulation. Genes which received a high fitness score, and thus were ranked in high positions in our list, were predicted to be potential regulators of FOXP3.

Any threshold separating high and low ranked genes would be somewhat arbitrary. However, plotting the fitness score against the model number (Fig. 1b), we remarked that two regimes can be identified, separated by a “knee” occurring after the first few hundred models. The first regime corresponded to higher fitness values, and the fitness value decreased steeply with model number. The second regime corresponded to lower fitness values, and it decreased almost linearly with the model number, until a final kink where fitness went to zero.

The upper part of the fitness scores, before the knee, corresponded to fitness values ranging from 62.14 to 45.78. These high fitness models included 176 models, which corresponded to approximately 2.5% of all models. We considered only these models as having a fitness high enough to represent a potential regulator of FOXP3 worth investigating further. The list of these 176 transcripts is reported in the Supplementary Material as Tab. S1. These 176 transcripts corresponded to 161 genes, since in 15 cases two transcripts corresponded to the same gene.

However, testing all of these 161 genes *in vitro* is practically impossible, due to the intrinsic limitations of available resources, and in practice we could test only five of these regulations *in vitro*. We thus needed to select five such promising regulatory genes for further laboratory experimental validation. Since the fitness scores of these genes were very close, we needed to choose which one to test based on other criteria, knowing that any criterion to select potential regulators will be somewhat arbitrary, but with the goal of selecting genes that were both likely to be regulators of FOXP3, and of potential biological relevance.

We thus performed this selection of five genes potentially regulating FOXP3 to be tested *in vitro* by means of the following considerations. First, transcripts were excluded when corresponding to more than one Entrez gene ID (more than one gene name due to the shared transcripts among genes) based on the HG-U133 plus 2.0 array annotation file (http://www.affymetrix.com/, version 24, 7 March 2008 [He et al., 2012]). Further, as already mentioned, the data of donor-2 were of higher quality (less affected by noise) w.r.t. those of donor-1, thus we only considered the data for donor-2 for selection purposes. Next, we required a reasonably clean dynamics of the time series signals, and a reasonable visually assessed dynamic behaviour. Furthermore, as mentioned we introduced a time-delay, and retained for each transcript only the model corresponding to the time-delay providing the highest fitness for each transcript. We also considered this delay information while performing our selection. We observed the time-difference between the peak of the candidate regulatory transcript and the peak of FOXP3 transcripts of donor-2. The reason behind this was that usually between the peak in mRNA production for a regulating transcript, and the peak of mRNA production of the regulated transcript, there usually need to be a certain amount of time which we considered to be reasonably between about 20 to 60 minutes to translate the mRNA of the first transcript into protein, which needs the time to bind to the promoter of the target transcript (FOXP3) and regulate its expression.

Having in these ways lowered down the 161 genes to a pull of 20, we performed a final selection on the basis of biological significance (gene ontology, based on http://amigo.geneontology.org/amigo, and protein location inside/outside the nucleus, based on https://www.uniprot.org/). On one hand, we preferred genes related to transcription and located in the nucleus, since the part of the regulatory pathways of FOXP3 which is largely unknown is predominantly the one occurring in the nucleus (see e.g. Lal and Bromberg [2009]). On these criteria, we selected 3 genes: NRBF2, NCOA7 and RNF12. On the other hand, we also did not want to limit ourselves to these predefined hypothesized functions and locations, so for the remaining two slots we selected genes representing different functions and with proteins located outside the nucleus, namely PDE4D and MAP1LC3B.

Thus based on all the criteria above we selected 5 regulatory genes for subsequent *in vitro* test, namely NCOA7, NRBF2, RNF12, PDE4D and MAP1LC3B.

#### 1.5 Software Availability

The codes developed in MATLAB which implement our computational approach and perform the described ranking of transcripts are available online at https://github.com/StefanoMagni/ModellingRegulatorsFOXP3.

**Table S1:**
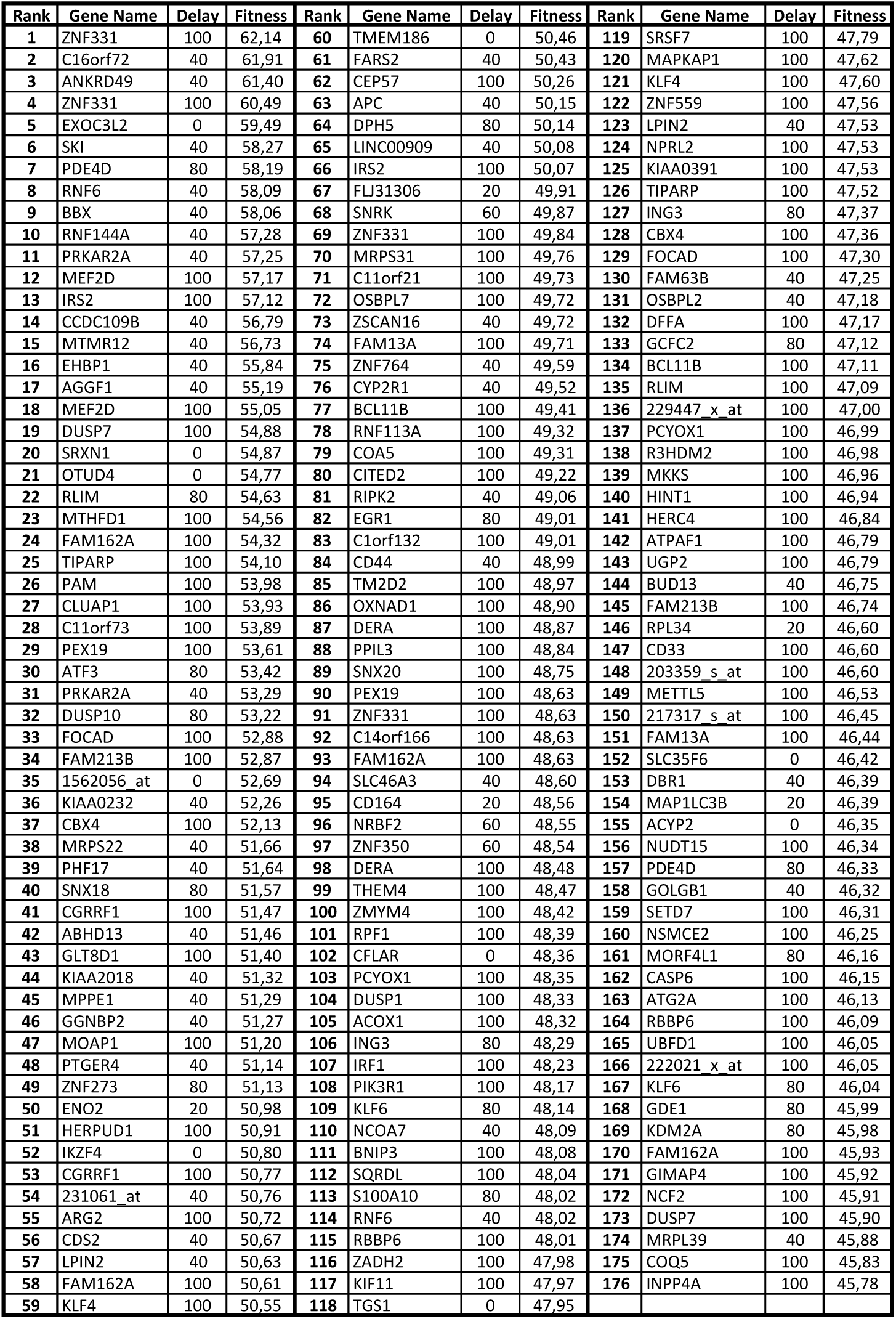
Highest ranked 176 transcripts, corresponding to 161 genes, according to our approach. First column: ranking, in descending order based on fitness scores. Second column: name of the gene, or Affymetrix probe-ID (a unique identifier) where multiple probe-IDs were associated to one name. Third column: delay (measured in minutes) corresponding to the model which received the highest fitness for that transcript as a regulator of FOXP3, for the second donor. Fourth column: actual value of the fitness score defined in Eq. (2), for that model (in percentage).

### 2 EXPERIMENTAL SUBJECTS, MODEL, METHODS DETAILS

#### 2.1 Regulatory T Cell Sorting and Culture

Informed consent was obtained from healthy blood donors through the Red Cross Luxembourg (RCL) and study procedures were approved by the ethic committee of the RCL. Buffy coats from healthy male donors of unknown age were provided by the RCL. Isolation of human Tregs was performed using similar methods as we described elsewhere [Danileviciute et al., 2019]. The RosetteSepac Human CD4+ T cell Enrichment Cocktail (15062, Stemcell) was added to undiluted blood at a concentration of 50 *µ*l/ml and incubated for 30 min at 4 *°*C. The blood was then diluted 2 times with FACS buffer (PBS + 2% FBS) and the CD4+ cells were isolated by gradient centrifugation at 1200 *g* for 20 min, using Lympoprep (07801, StemCell) and SepMateac-50 tubes (85450, Stemcell). Primary natural regulatory T cells (CD4+CD25^*high*^CD127^*low*^) were then sorted on a BD Aria III Flow cytometry cell sorter (BD Biosciences). CD4+ T cells were stained for 30 min with mouse monoclonal [RPA-T4] anti-human CD4 FITC (555346, BD Biosciences) (dilution 1:20), mouse monoclonal [M-A251] anti-human CD25 APC (555434, BD Biosciences) (dilution 1:20), mouse monoclonal [HIL-7R-M21] anti-human CD127 V450 (560823, BD Biosciences) (dilution 1:20) 4 *°*C followed by 2 washing steps with FACS buffer (250 *g*, 10 min).

Sorted Treg were cultured in IMDM (21980-032, ThermoFisher Scientific) complete medium, supplemented with 10% heat-inactivated (56 *°*C, 45 min) fetal bovine serum (FBS) (10500-064, ThermoFisher Scientific), 1x Penicillin+Streptomycin (15070-063, ThermoFisher Scientific), 1x MEM nonessential amino acids (M7145, Sigma-Aldrich) and 1x *β*-mercaptoethanol (21985-023, ThermoFisher Scientific) at 37 *°*C, 7.5% CO_2_. The medium was further supplemented with 100 U/ml of recombinant human IL2 (2238131, Novartis) for the culture of Tregs. For a maximum duration of 6 weeks, the Tregs were restimulated every 7 days with irradiated Epstein Barr virus (EBV) (strain B95-8, VR-1491, ATCC) transformed B cells (EBV-B cells), at a 1:1 ratio to expand and maintain the culture. A RS2000 X-Ray Biological Irradiator was used to irradiate the EBV-B cells (Rad Source Technologies) for 30 minutes with a total of 90 Gy. On a regular basis, the Tregs were characterized by Flow Cytometry for their expression of CD4, CD25, FOXP3 and Helios and discarded if the expression of FOXP3 and/or Helios was apparently decreased.

#### 2.2 Flow Cytometry for Treg Characterisation

Extracellular markers were stained in FACS buffer for 30 min at 4 *°*C, followed by 3 washing steps (250 *g*, 10 min). Fixation, permeabilization and staining of intracellular markers was performed using the True-Nuclear Transcription Factor Buffer Set (424401, BioLegend) and following the manufuacturers instructions. The antibodies used for the characterisation are in the the following: mouse monoclonal [RPA-T4] anti-human CD4 BUV395 (564724, BD Biosciences) (dilution 1:100), mouse monoclonal [M-A251] anti-human CD25 FITC (555431, BD Biosciences) (dilution 1:100), mouse monoclonal [22F6] anti-human Helios Pacific blue (137220, BioLegend) (dilution 1:100), mouse monoclonal [206D] anti-human FOXP3 Alexa Fluor 647 (320114, BioLegend) (dilution 1:20). LIVE/DEAD® Fixable Near-IR Dead Cell Stain (L10119, ThermoFisher Scientific) (dilution 1:500).

#### 2.3 Treg siRNA Knockdown and Stimulation

The P3 Primary Cell 4D-Nucleofector X Kit L (V4XP-3024, Lonza) was used for the knock-down of the targeted genes, using 90 *µ*l P3 Primary cell solution and 100 pmol of corresponding si_RNA (resuspended in 10 *µ*l RNAse-free H2O): si_Non-Specific (si_NS or si_CTL) (sc-37007, Santa Cruz), si_NRBF2 (SI00139118, Qiagen), si_NCOA7 (SI02649668, Qiagen), si_MAP1LC3B (SI04200735, Qiagen), si_PDE4D (SI05587666, Qi-agen), si_RNF14 (SI00113582, Qiagen). The Amaxa 4D-Nucleofector X System (Lonza) was used to perform the electroporation and siRNA transfection according to the manufacturers recommended program for primary human T cells. After transfection, the Tregs were transferred into a 12-well plate with pre-warmed complete medium, supplemented with 100 U/ml IL-2, and kept at 37 *°*C for 24 h before being stimulated with 25 *µ*l/ml of soluble antibodies (Immunocult Human CD3/CD28 T Cell Activator) (10971, Stemcell) in a 24-well plate for 5, 10 and 15 h.

#### 2.4 RNA Extraction

RNA was extracted using the RNeasy Mini Kit (74106, Qiagen), following the manufacturers instructions and including the digestion of genomic DNA with DNAse I (79254, Qiagen). The cells were lysed in RLT buffer (Qiagen), supplemented with 1% beta-Mercaptoethanol (63689, Sigma-Aldrich). The NanoDrop 2000c Spectrophotometer (Thermo Fisher Scientific) was used to measure the RNA concentration and its quality was checked by assessing the RNA integrity number (RIN) using the Agilent RNA 6000 Nano kit (5067-1511, Agilent) and the Agilent 2100 Bioanalyzer Automated Analysis System (Agilent), according to the manufacturers protocol. Only the samples with RIN of 8 or higher were considered for further analysis.

#### 2.5 cDNA Synthesis

A maximum of 500 ng of RNA was used for human cDNA synthesis, using the Superscipt IV First Strand Synthesis System (18091050, ThermoFisher Scientific) and following the manufacturers instructions. For the first step the mastermix contained following components (per sample): 0.5 *µ*l of 50 *µ*M Oligo(dT)20 primers (18418020, ThermoFisher Scientific), 0.5 *µ*l of 0.09 U*µ*l Random Primers (48190011, ThermoFisher Scientific), 1 *µ*l of 10 mM dNTP mix (18427013, ThermoFisher Scientific) and RNAse free water for a final volume of 13 *µ*l in 0.2 ml PCR Tube Strips (732-0098, Eppendorf). The reaction tubes were transferred into a C1000 Touch Thermal Cycler (Bio-Rad) or UNO96 HPL Thermal Cycler (VVR) and subjected to the following program: 5 min at 65 *°*C, followed by 2 min at 4 *°*C. For the second step, the reaction mix was supplemented with 40 U RNaseOUT Recombinant Ribonuclease Inhibitor (10777019, ThermoFisher Scientific), 200 U SuperScript IV Reverse Transcriptase (18090050, ThermoFisher Scientific), a final concentration of 5 mM Dithiothreitol (DTT) (70726, ThermoFisher Scientific) and 1x SSIV in a total reaction volume of 20 *µ*l. The thermocycler program for the second step was the following: 50 *°*C for 10 min, then 80 *°*C for 10 min and 4 *°*C until further usage. The obtained cDNA was then 5x diluted with nuclease-free water to a final volume of 100 *µ*l.

#### 2.6 Quantitative Real-Time PCR (qPCR)

The reaction mix per sample for quantitative realrime PCR (qPCR) contained: 5 *µ*l of the LightCycler 480 SYBR Green I Master Mix (04707516001, Roche), 2.5 *µ*l cDNA and 2.5 *µ*l primers in a total reaction volume of 10 *µ*l. The reaction was performed in a Light-Cycler 480 (384) RT-PCR platform (LightCycler 480 (384), Roche), using the LightCycler 480 Multiwell 384-well plates (04729749 001, Roche) and LC 480 Sealing Foil (04729757001, Roche). The temperature program for qPCR was the following: 95 *°*C for 5 min; 45 cycles of (55 *°*C for 10 s, 72 *°*C for 20 s, 95 °C for 10 sec); meltingcurve (65-97 *°*C). The results were analysed with the LightCycler 480 SW 1.5 software. Primers used for qPCR: RPS9 (QT00233989, Qiagen) as a reference gene, NRBF2 (QT00061936, Qiagen), NCOA7 (QT00033922, Qiagen), FOXP3 (QT00048286, Qiagen), PDE4D (QT00019586, Qiagen) and MAP1LC3B (QT00055069, Qiagen).

#### 2.7 Western Blotting

Proteins were separated in Novex WedgeWell 4-20% Tris-Glycine Gels (XPO4202Box, Invitrogen), using the Novex Tris-Glycine SDS Running buffer (LC2675-4, Invitrogen). The proteins were transferred (dry transfer) using an iBlot2 Gel Transfer Device (IB21001, Invitrogen) and iBlot2 PVDF stacks (IB24002, Invitrogen). After the transfer the membranes were blocked in 5% milk in PBS with 0.2% Tween20 (PBS-T) for 1 h at room temperature with gentle shaking before being incubated overnight at 4 *°*C with the primary antibodies: rabbit monoclonal [15H7L3] anti-human NRFB2 (702920, ThermoFisher Scientific) (dilution 1:5000), rabbit polyclonal [FL-335] GAPDH (sc-25778, Santa Cruz Biotechnology) (dilution 1:200), mouse monoclonal [206D] FOXP3 (320102, Biolegend) (dilution 1:100), diluted in 5% BSA in PBS-T with 0.025% sodium azide. The next day the membrane was washed three times for 10 min before and after incubation with secondary goat anti-rabbit HRP-coupled antibodies (172-1019, Bio-Rad). The proteins were detected using the Amersham ECL Prime Western Blotting Detection Reagent (RPN2232, GE Healthcare Life Sciences) and visualized on the ECL Chemocam Imager (INTAS). If needed, the contrast and brightness of the obtained whole picture was adjusted using the Fiji ImageJ software.

## KEY RESOURCES TABLE

**Table S2:** Key Resources. Materials employed for the experimental methods described in the text.

| REAGENT or RESOURCE | SOURCE | IDENTIFIER |
| --- | --- | --- |
| **Antibodies** |  |  |
| mouse monoclonal [RPA-T4] anti-human CD4 FITC (dilution 1:20) | BD Biosciences | Cat# 555346 |
| mouse monoclonal [M-A251] anti-human CD25 APC (dilution 1:20) | BD Biosciences | Cat# 555434 |
| mouse monoclonal [HIL-7R-M21] anti-human CD127 V450 (dilution 1:20) | BD Biosciences | Cat# 560823 |
| mouse monoclonal [RPA-T4] anti-human CD4 BUV395 (dilution 1:100) | BD Biosciences | Cat# 564724 |
| mouse monoclonal [M-A251] anti-human CD25 FITC (dilution 1:100) | BD Biosciences | Cat# 555431 |
| mouse monoclonal [22F6] anti-human Helios Pacific blue (dilution 1:100) | BioLegend | Cat# 137220 |
| mouse monoclonal [206D] anti-human FOXP3 Alexa Fluor 647 (dilution 1:20) | BioLegend | Cat# 320114 |
| LIVE/DEAD® Fixable Near-IR Dead Cell Stain (dilution 1:500) | ThermoFisher Scientific | Cat# L10119 |
| rabbit monoclonal [15H7L3] anti-human NRFB2 (dilution 1:5000) | ThermoFisher Scientific | Cat# 702920 |
| Purified anti-human FOXP3 Antibody (dilution 1:100) | BioLegend | Cat# 320102 |
| rabbit polyclonal [FL-335] GAPDH (dilution 1:200) | Santa Cruz Biotechnology | Cat# sc-25778 |
| goat anti-rabbit HRP-coupled antibodies | Bio-Rad | Cat# 172-1019 |
| Immunocult™ Human CD3/CD28 T Cell Activator | StemCell | Cat# 10971 |
| **Bacterial and Virus Strains** |  |  |
| Epstein-Barr virus, strain B95-8 | ATCC | Cat# VR-1491 |
| **Biological Samples** |  |  |
| Regulatory T cells, isolated from buffy coats from healthy male adults of unknown age. | Center for Blood Transfusions, Red Cross Luxembourg | http://www.croix-rouge.lu/de/locations/centre-de-transfusion-sanguine/ |
| **Chemicals, Peptides, and Recombinant Proteins** |  |  |
| Recombinant human Interleukin-2 | Novartis | Cat# 2238131 |
| **Critical Commercial Assays** |  |  |
| RosetteSep™ Human CD4+ T cell Enrichment Cocktail | StemCell | Cat# 15062 |
| SepMate™-50 tubes | StemCell | Cat# 85450 |
| Lymphoprep | StemCell | Cat# 07801 |
| True-Nuclear Transcription Factor Buffer Set | BioLegend | Cat# 424401 |
| P3 Primary Cell 4D-Nucleofector X Kit L | Lonza | Cat# V4XP-3024 |
| RNeasy Mini Kit | Qiagen | Cat# 74106 |
| Superscript™ IV First Strand Synthesis System | ThermoFisher Scientific | Cat# 18091050 |
| LightCycler 480 SYBR Green I Master Mix | Roche | Cat# 04707516001 |
| Novex™ WedgeWell 4-20% Tris-Glycine Gels | Invitrogen | Cat# XPO4202Box |
| Novex™ Tris-Glycine SDS Running buffer | Invitrogen | Cat# LC2675-4 |
| Amersham ECL Prime Western Blotting Detection Reagent | GE Healthcare Life Sciences | Cat# RPN2232 |
| **Deposited Data** |  |  |
| **Experimental Models: Cell Lines** |  |  |
| **Experimental Models: Organisms/Strains** |  |  |
| **Oligonucleotides** |  |  |
| Control siRNA | Santa Cruz | Cat# sc-37007 |
| Hs_NRBF2_3 FlexiTube siRNA | Qiagen | Cat# SI00139118 |
| Hs_NCOA7_7 FlexiTube siRNA | Qiagen | Cat# SI02649668 |
| Hs_MAP1LC3B_8 FlexiTube siRNA | Qiagen | Cat# SI04200735 |
| Hs_PDE4D_9 FlexiTube siRNA | Qiagen | Cat# SI05587666 |
| Hs_RNF12_1 FlexiTube siRNA | Qiagen | Cat# SI00113582 |
| Hs_RPS9_1_SG QuantiTect Primer Assay | Qiagen | Cat# QT00233989 |
| Hs_NRBF2_1_SG QuantiTect Primer Assay | Qiagen | Cat# QT00061936 |
| Hs_NCOA7_1_SG QuantiTect Primer Assay | Qiagen | Cat# QT00033922 |
| Hs_FOXP3_1_SG QuantiTect Primer Assay | Qiagen | Cat# QT00048286 |
| Hs_CTLA4_2_SG QuantiTect Primer Assay | Qiagen | Cat# QT01670550 |
| Hs_CD4_1_SG QuantiTect Primer Assay | Qiagen | Cat# QT00005264 |
| Hs_MAP1LC3B_1_SG QuantiTect Primer Assay | Qiagen | Cat# QT00055069 |
| Hs_PDE4D_1_SG QuantiTect Primer Assay | Qiagen | Cat# QT00019586 |
| Hs_RLIM_1_SG QuantiTect Primer Assay | Qiagen | Cat# QT00020524 |
| **Recombinant DNA** |  |  |
| **Software and Algorithms** |  |  |
| ImageJ | Fiji | https://imagej.net/Fiji/Downloads |
| **Other** |  |  |

